# Photochemical pre-bleaching of formalin-fixed archival prostate tissues significantly reduces autofluorescence to facilitate multiplex immunofluorescence staining

**DOI:** 10.1101/2021.11.09.467916

**Authors:** Alan K. Meeker, Christopher M. Heaphy, Christine M. Davis, Sujayita Roy, Elizabeth A. Platz

## Abstract

The characterization of tissues using multiple different primary antibodies detected by secondary antibodies, each possessing a different colored fluorophore (multiplex immunofluorescence), is a powerful technique but often impaired by endogenous autofluorescence present in the specimen. Our current research involves the use of multiplex immunofluorescence to identify specific cell phenotypes within the tumor microenvironment in archival formalin-fixed paraffin-embedded human prostate cancer tissue specimens. These specimens frequently possess high levels of autofluorescence, in part due to the biological age of the tissues and long storage times. This autofluorescence interferes with and, in the worst cases, completely obscures the desired immunofluorescent signals, thus impeding analyses by decreasing signal-to-noise. Here, we demonstrate that a recently published protocol for photochemical bleaching significantly decreases autofluorescence (80% average decrease of the brightest autofluorescent signals), across the visible spectrum, in fixed, archival prostate tissue specimens from aged men, that have been sectioned onto glass slides and stored for several months. Importantly, the method is compatible with subsequent immunofluorescence staining and yields markedly improved signal-to-noise. Inclusion of this method should facilitate studies employing multiplex immunofluorescence in sections cut from archival fixed human prostate tissues.

## Introduction

Immunofluorescence (IF) applied to tissue specimens is a powerful technique, allowing for the sensitive and specific detection of proteins of interest within their histomorphologic context (1,2). IF can be used to highlight particular tissue structures (e.g. blood vessels), as well as determine cell identities (e.g. T cells, cancer cells, stromal cells) or cellular phenotypes (e.g. proliferative) based on their expression of specific protein biomarkers. Computer-assisted image analysis of digital microscopy images of IF-stained tissues can provide quantitative measurements, such as enumeration of specific cells, the determination of the relative expression levels of specific proteins, or the spatial localization of specific cell populations. The advent of a broad range of fluorescent dyes and specialized IF staining protocols enables multiplex, multicolor IF, thereby providing additional layers of spatial information from a single archival tissue specimen (2). However, in contrast to other staining techniques (e.g. chromogenic immunohistochemistry), cell and tissue-based autofluorescence is a problem commonly encountered in IF. In the context of IF, autofluorescence refers to undesired background fluorescence that decreases signal-to-noise, thus confounding target visualization, image analysis, and, in extreme cases, making it impossible to detect the specific IF signals altogether (3,4).

In archival tissue specimens, autofluorescence originates from a wide variety of endogenous sources. These sources include the presence of common structural proteins (e.g. collagen and elastin), as well as proteins containing disproportionately high amounts of aromatic amino acids, nicotinamide compounds (NAD and NAD(P)H), oxidized flavins, melanin, or folic acid. Additionally, cellular organelles such as mitochondria and lysosomes, as well as intact red blood cells and their residual breakdown products can increase autofluorescence (5–10). Autofluorescence can also be caused by the process of tissue fixation, particularly in tissues fixed with the routinely used aldehyde-based fixatives formaldehyde and glutaraldehyde that react with amines, thereby generating fluorescent byproducts. Further exacerbating the problem, these autofluorescent byproducts can increase over time during storage, and can increase through heating processes, such as when cut sections from formalin-fixed paraffin-embedded (FFPE) tissue blocks are baked onto microscope slides to promote their adherence (11,12). These problems are of particular concern when conducting retrospective translational studies utilizing archival tissue samples, which typically consist of FFPE specimens stored for years to decades. FFPE tissue sections also require aqueous heat pretreatment (so-called “epitope retrieval”) prior to the conduct of IF staining, which may increase autofluorescence. Finally, sources of autofluorescence can accumulate over time in vivo as the organism ages. These sources include the so-called “age-pigment” lipofuscin, as well as advanced glycation end products (AGEs), the result of glycation reactions between sugars and proteins or lipids (6,13,14). The autofluorescence observed in FFPE tissues typically covers a broad spectrum of colors in the visible range, from blue to red. Thus, this autofluorescence can potentially interfere with any of the fluorescent dyes commonly used to label antibodies for conducting IF, and therefore, is particularly problematic when attempting to conduct multiplex, multicolor IF.

In order to minimize autofluorescence, tissues can be fixed with alternative, non-aldehyde fixatives, or preserved by freezing rather than fixation. However, these alternative methods are often unavailable (e.g. if using archival, fixed tissues collected for clinical purposes), and they do not prevent native autofluorescence that is not due to fixation. Several approaches have thus been attempted with the goal of reducing or eliminating autofluorescence, including treatment of tissue sections with chemical fluorescence quenching agents or dyes (e.g. sodium borohydride, trypan blue, eriochrome black T, ammonium chloride and copper sulfate), or with lipophilic dyes (e.g. Sudan black). Despite many attempts, results with such agents have been inconsistent and dependent upon a number of variables, including tissue type, fixation, and storage details. Further, these agents often i) only target a subset of the many autofluorescent species present, ii) may shift the autofluorescence spectrum rather than reduce or eliminate it, iii) may require cumbersome, labor-intensive protocols, iv) can damage the tissue or cause detachment of the tissue from the slide, or v) can even result in *increased* autofluorescence of some tissue components, as well as decrease the desired IF signals (15–18). In addition to chemical treatments, pre-bleaching of tissue sections has been attempted via exposure of the tissue to either UV or visible light. These methods may require specialized equipment not typically present in microscopy laboratories, and, similar to chemical agents, have resulted in mixed and inconsistent results (17–19).

For single or dual-target IF staining studies, optimal results often are obtained by utilizing fluorescent dyes that emit light in the near infrared (NIR) part of the spectrum, where tissue autofluorescence tends to be less intense. However, drawbacks of this approach, include the inability of most individuals to directly visualize such signals by eye through the microscope, thereby requiring the aid of a digital camera, and the ability to use only one or two antibodies within the relatively narrow NIR portion of the spectrum, thus disallowing higher-order multiplex IF studies. When conducting IF staining studies, other solutions to the problem of tissue autofluorescence, include the use of specialized microscopy platforms, such as confocal microscopy or multispectral imaging approaches, both of which require specialized equipment, expertise, and computation for image collection and downstream image deconvolution, and are relatively low throughput, thus hampering the analysis of large numbers of samples (20–22).

Our laboratory conducts molecular patho-epidemiologic studies in human prostate cancer, thus requiring the conduct of multiplex IF staining on hundreds of archival FFPE prostate specimens sourced from large clinical and epidemiologic cohorts. IF staining studies using FFPE human prostate tissues are typically plagued by a wide variety of autofluorescent species, including endogenous sources described above, and formalin fixation-induced fluorescent species exacerbated by long storage times in pathology archives. In addition, these prostate tissues were resected from older men, and thus likely have also accumulated substantial amounts of age-related autofluorescent species such as lipofuscin, advanced glycation products, and intra-ductal entities termed corpora amylacea (23,24).

Recently, Du et al. described a method for reducing autofluorescence in FFPE tissue specimens by chemical bleaching with hydrogen peroxide combined with exposure to an intense visible light source (25). Here, we tested the effectiveness of this method, as well as its compatibility with subsequent immunofluorescent staining, in archival human FFPE prostate tissue sections. These specimens frequently possess high levels of autofluorescence, in part due to the biological age of the tissues and long storage times that interfere with and, in the worst cases, can completely obscure the desired immunofluorescence signals, thus impeding or preventing analyses by decreasing signal-to-noise. Here, we demonstrate that the method markedly reduces background autofluorescence, across the visible spectrum, in archival, fixed human FFPE prostate tissue sections. This method is compatible with downstream IF staining, and greatly facilitates multiplex IF by increasing the signal-to-noise of the desired immunofluorescence signals, at multiple wavelengths.

## Materials and Methods

### Tissue samples

For this study, we utilized serial, adjacent, four-micrometer thick sections cut from de-identified archival human prostate FFPE prostatectomy specimens from men who underwent surgical resection at least five years prior to use in this study. Cut sections were applied to charged microscope slides and the slides stored at −20 degrees Celsius for several months prior to use in this study. These samples were chosen as representative of archival FFPE prostatectomy tissue samples used in tissue-based, retrospective cohort studies. These tissues typically have significant background fluorescence due, in part, to the fact that prostate cancer is primarily a disease affecting older men, and because autofluorescence in fixed tissues is thought to increase over time during paraffin block storage, and during storage of tissue sections cut to slides. Archival FFPE blocks in large cohort studies of prostate cancer are typically many years old, particularly when the study focus is on comparing tissue-based measurements with post-surgical clinical outcomes, as prostate cancer is a slowly progressing disease.

### Photochemical bleaching of autofluorescence

We utilized the protocol of Du, et al. (25), as described. Briefly, FFPE tissue sections were rehydrated by immersion through xylenes, 100% ethanol, 95% ethanol, 70% ethanol, and then in distilled water. Heat-induced antigen retrieval was performed in a pH 6.0 sodium citrate buffer heated to 100 degrees Celsius in a steamer for 15 minutes, followed by cooling to room temperature. Slides were then transferred to phosphate buffered saline (PBS). Slides were immersed in 30 mL of alkaline hydrogen peroxide solution in clear plastic petri dishes and exposed to white light by sandwiching the immersed slides between two light-emitting diode (LED) panels, as described (25). The slides were incubated for 45 minutes, after which time the bleaching solution was discarded and replaced by freshly-made bleaching solution, and the 45 minute light exposure was repeated. Slides were then washed in PBS and immersed in PBS until used for immunofluorescence. Unbleached control slides were treated identically as bleached slides, except that all bleaching steps, including peroxide solution and LED light exposures were omitted. Instead, control slides remained immersed in PBS at room temperature in room light while the remaining slides were subjected to bleaching.

### Immunofluorescence

Following photochemical bleaching, immunofluorescent staining was performed using standard protocols. Briefly, rabbit-derived anti-CD4 primary antibody (Sino Biological, Cat# 10400-R), diluted 1:200 in antibody dilution buffer (Ventana, Cat# ADB240), was applied to the slides and incubated for 45 minutes at room temperature. Slides were then washed three times in Tris-buffered saline+Tween-20 (TBST). 200 uL of anti-rabbit secondary antibody (Leica Cat# PV6119, diluted 1:125 in PBS) was added to each slide and incubated 30 minutes at room temperature. Slides were then washed three times in TBST. 150 uL of Opal520 (Akoya Biosciences, Cat# FP1487001KT) working solution/fluorophore was applied to each slide and incubated 10 minutes at room temperature followed by three TBST washes. Blocking buffer (Dako, Cat# S2003) was then applied for 10 minutes at room temperature, followed by rinsing with TBST. Mouse-derived anti-CD8 primary antibody (Dako, Cat# M7103), diluted 1:100 in Antibody Dilution Buffer, was then applied to the slides and incubated for 45 minutes at room temperature. Slides were washed three times in Tris-buffered saline+Tween-20 (TBST). 200 uL of anti-mouse secondary antibody (Leica Cat# PV6114, diluted 1:125 in PBS) was added to each slide and incubated 30 minutes at room temperature. Slides were then washed three times in TBST. 150 uL of Opal570 (Akoya Biosciences, Cat# FP1487001KT) working solution/fluorophore was applied to each slide and incubated 10 minutes at room temperature followed by three TBST washes. In some sets of slides, nuclei were then counterstained with DAPI. Cover slips were mounted with ProLong Gold mounting medium (Thermo-Fisher Cat# P36930), and slides were then stored at 4 degrees Celsius in the dark until viewing and imaging.

### Image capture and analysis

Slides were imaged using a Nikon 50i epifluorescence microscope equipped with an X-Cite illumination system (EXFO Photonics Solutions), 10X or 20X primary objectives, and fluorescence excitation/emission filter sets appropriate for each separate fluorophore (CD4-green channel, CD8-red channel, DAPI-blue channel). The same system and filter sets were used to capture images of tissue autofluorescence in unstained slides, either unbleached or bleached. Individual 12-bit greyscale images were captured using a Photometrics CoolsnapEZ digital camera and stored as uncompressed TIFF files through Nikon NIS-Elements imaging software. Quantification of image intensity signals (total intensity for all pixels in the entire image; as well as minimum, maximum and mean intensity values) was performed by importing the uncompressed TIFF greyscale images into the open source image analysis program, ImageJ (https://imagej.nih.gov/ij/). 3D perspectival plots of image intensities were generated in NIS-Elements. When capturing images, exposure times for each of the separate color channels were kept constant across all slides within each comparison set (e.g. bleached and unbleached slides) during image collection. Likewise, all comparison sets of adjacent slides from the same case were processed and run together in the same bleaching runs and during IF staining.

## Results

We tested the effectiveness of a previously published tissue photochemical bleaching protocol (25) in reducing endogenous autofluorescence in archival FFPE human prostate tissues. We used serial four micrometer thick sections from paraffin blocks that were at least five years old, placed on charged microscope slides and stored in a −20 degree Celsius freezer for greater than three months prior to use. As described in Materials and Methods, the published protocol involves two cycles of immersion of the tissue slides in an alkaline hydrogen peroxide bleaching solution for 45 minutes while exposed to a bright white LED light source. As shown in **Figure 1**, we observed significant qualitative reductions in autofluorescence in the red and green channels when the same tissue sections were compared before and after bleaching. In order to quantify the amount of of autofluorescence reduction, digital images were captured for three different spectral regions commonly used in immunofluorescence studies (red, green, and aqua) from the same physical location within the adjacent bleached and unbleached tissue specimens. For each color channel, the camera exposure times were identical for both the bleached and unbleached specimens, and all slides within the same batch were run simultaneously. Each image was subsequently analyzed using the open source, Java-based image processing and analysis software, ImageJ. Pixel intensities in each image were analyzed to determine total, mean, minimum and maximum pixel signal intensities. As shown in **Figure 2**, mean, maximum and total autofluorescence intensities were markedly reduced, in all three colors assessed, in the bleached samples compared to the unbleached adjacent tissue sections. This reduction in autofluorescence is largely due to suppression of the most intense areas of autofluorescence, as reflected in the ~80% loss of maximum autofluorescence signals, in all three-color channels. Examples of the overall reduction of endogeneous autofluorescence by bleaching is provided by the 3D fluorescence intensity contour plots in **Figure 3**. To assess the durability of the observed reduction in autofluorescence we re-evaluated these same slides after one month of storage at four degrees Celsius and found no appreciable restoration of autofluorescence in the bleached tissue sections (data not shown).

**Figure 1.**
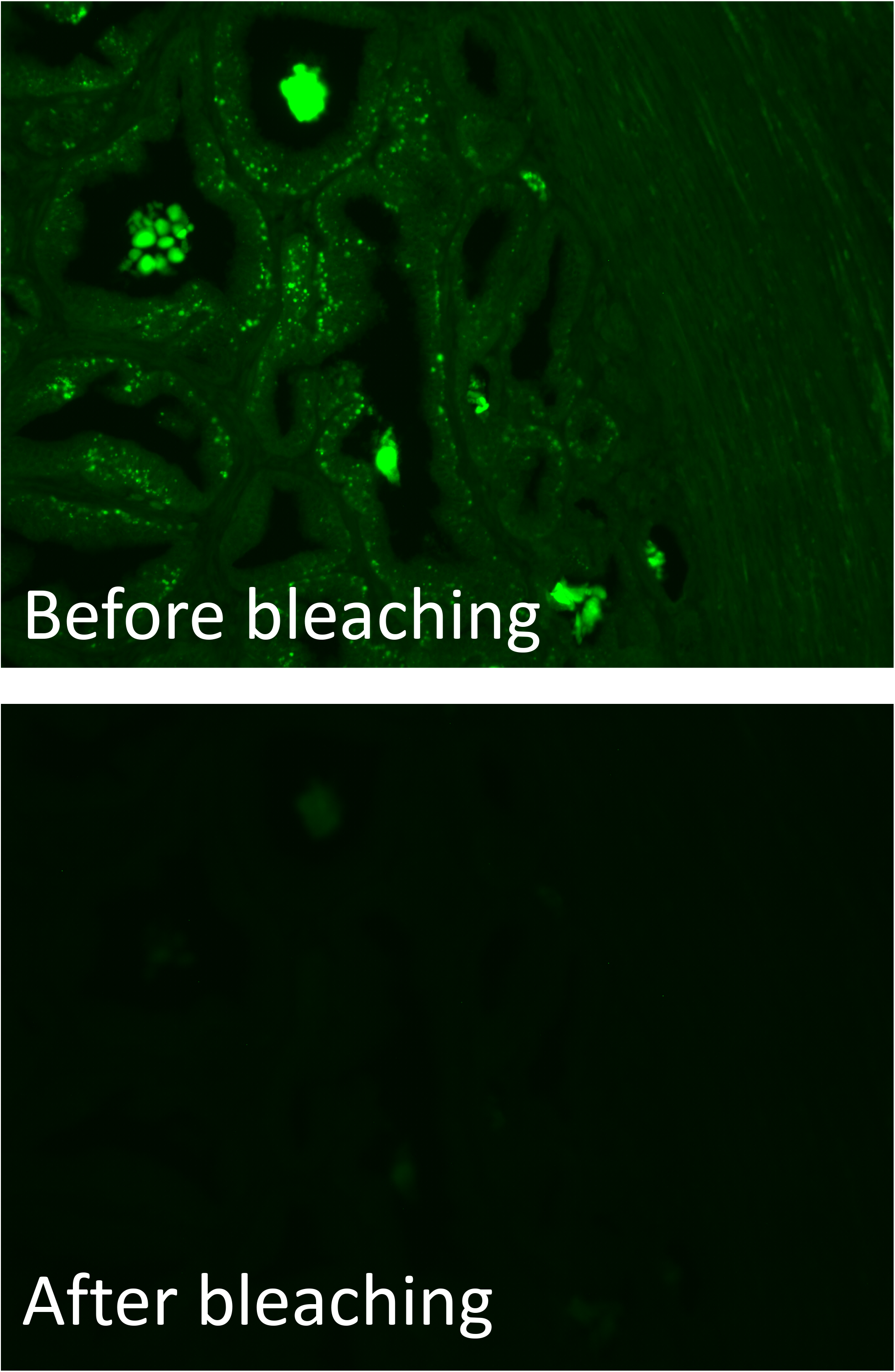
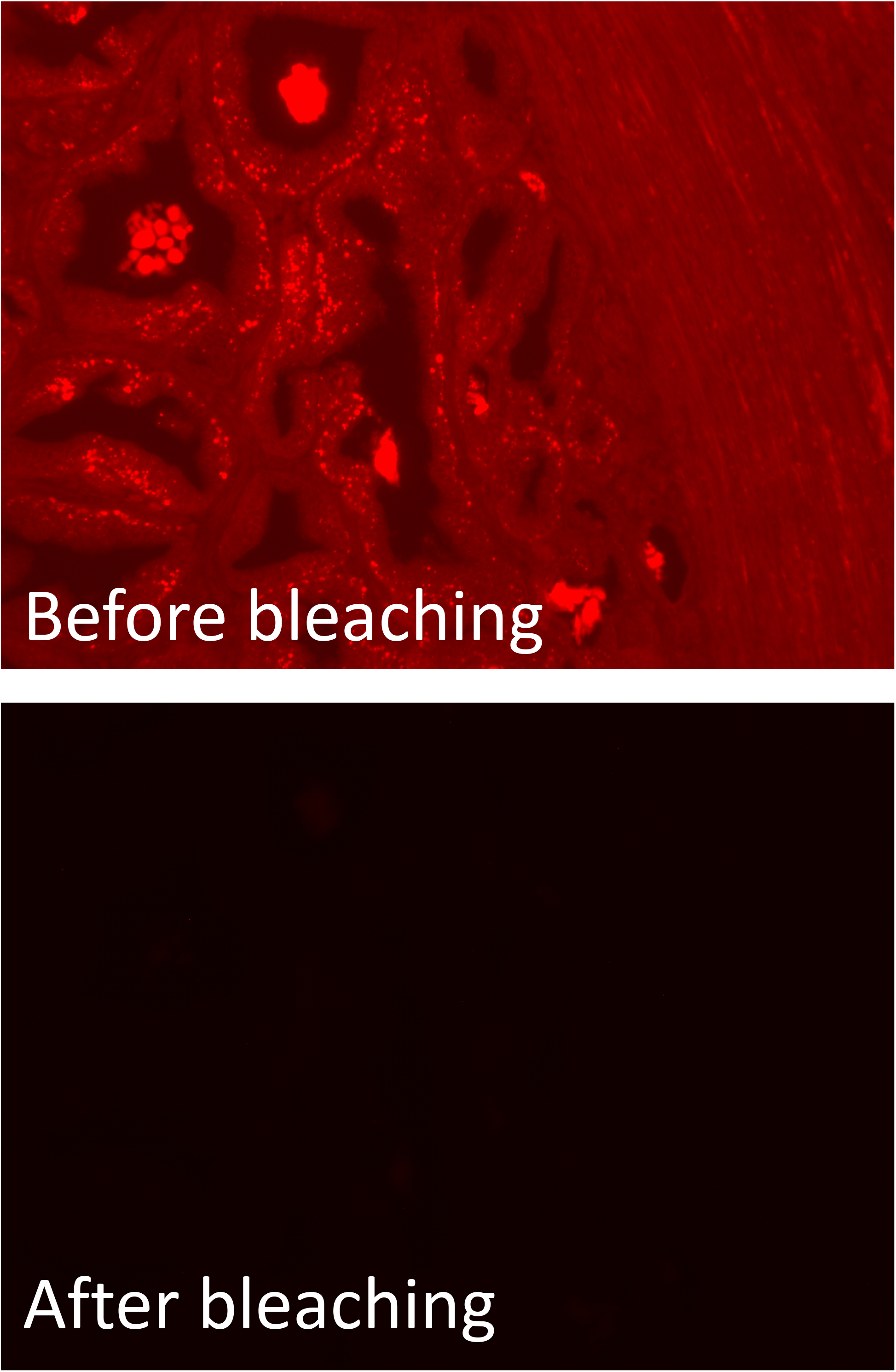
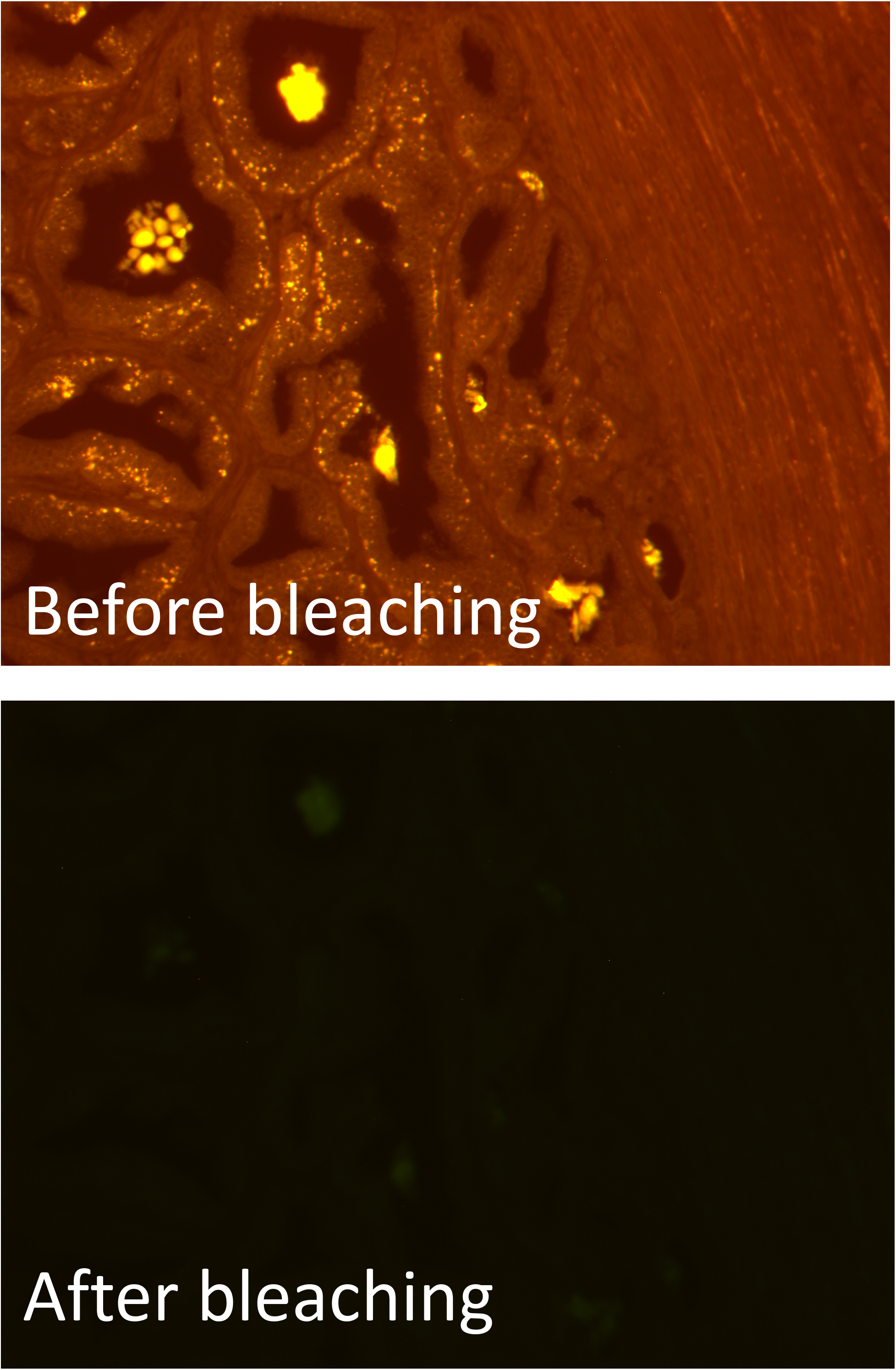
Photochemical bleaching markedly reduces autofluorescence in FFPE prostate tissue. Fluorescence images of an unstained section of FFPE prostate, imaged either before bleaching (upper image) or after bleaching (lower image). A) Green fluorescence channel; B) Red fluorescence channel; C) Combined images from green and red channel. For each color channel, exposure times used were identical for collecting images before bleaching and after bleaching. All images were captured using a 10X objective.

**Figure 2.**
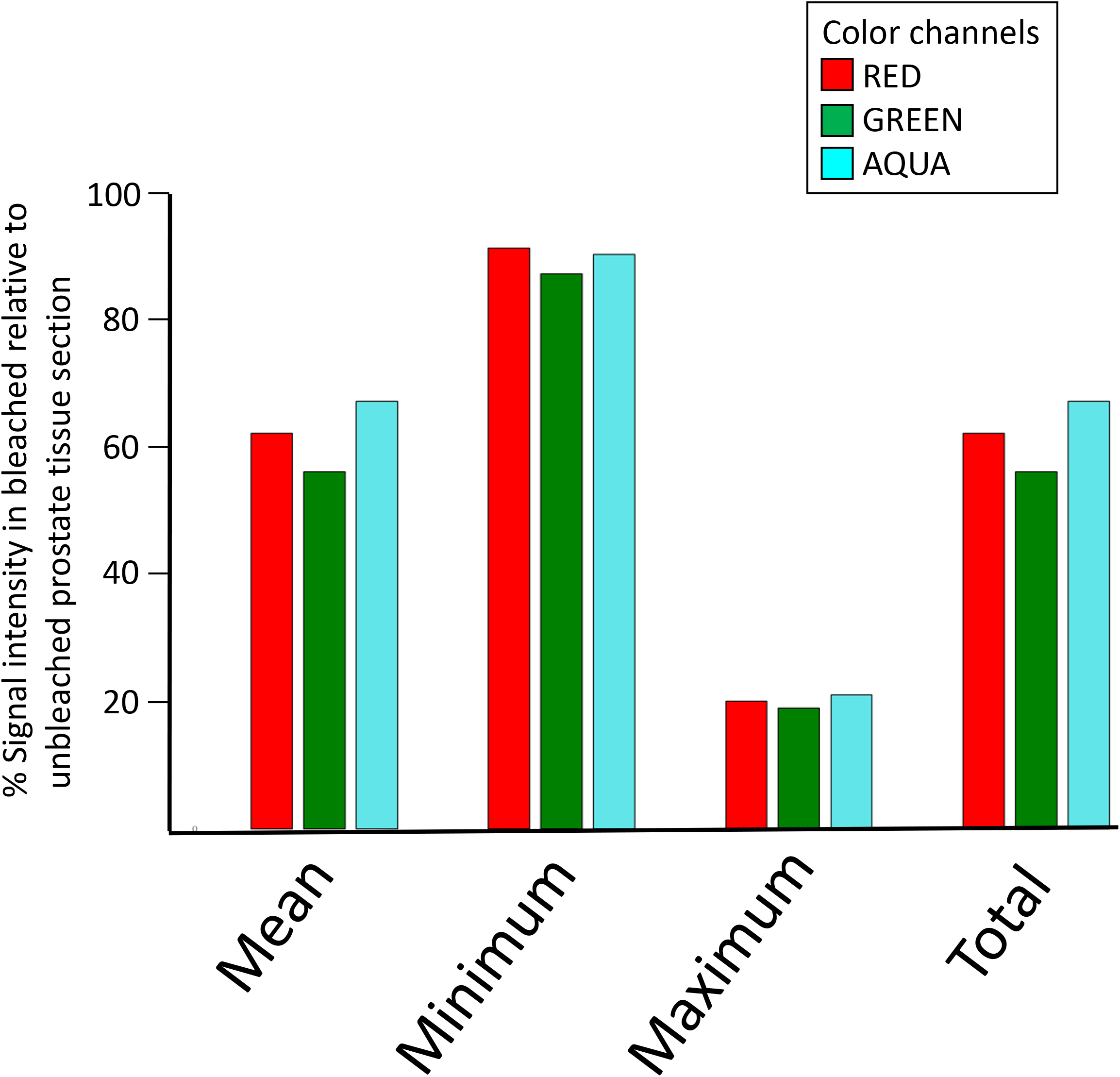
Quantification of the amount of autofluorescent reduction in FFPE prostate sections due to photochemical bleaching. Fluorescence intensity data from images of a bleached, unstained tissue sections of an FFPE prostate specimen (different specimen than used in Figure 1), in three different color channels (red, green and aqua). Values for mean, minimum, maximum, and total fluorescence intensities were calculated from whole 20X images and are plotted relative to the respective values calculated for an adjacent unbleached section. All images were captured from the same area of the specimen.

**Figure 3.**
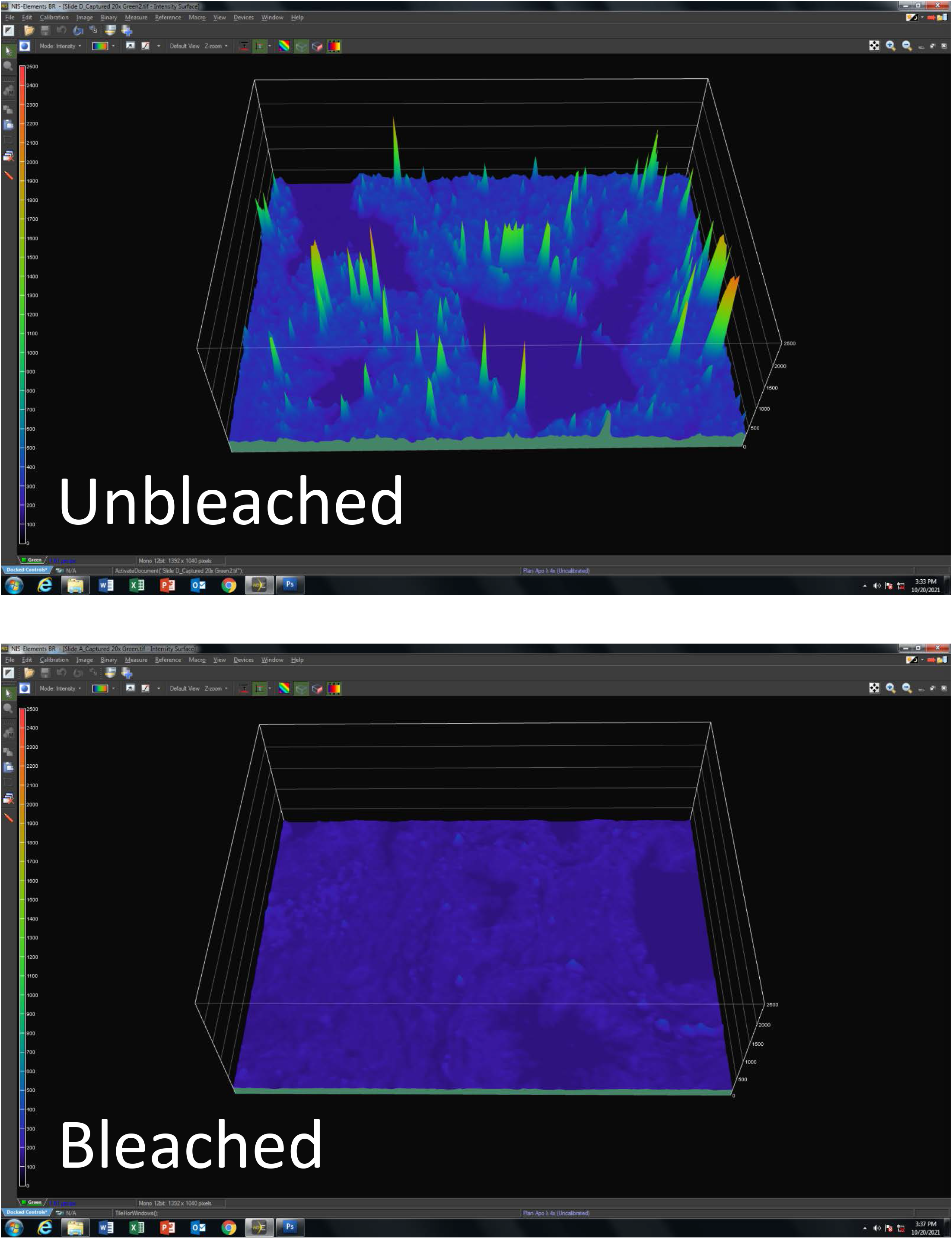
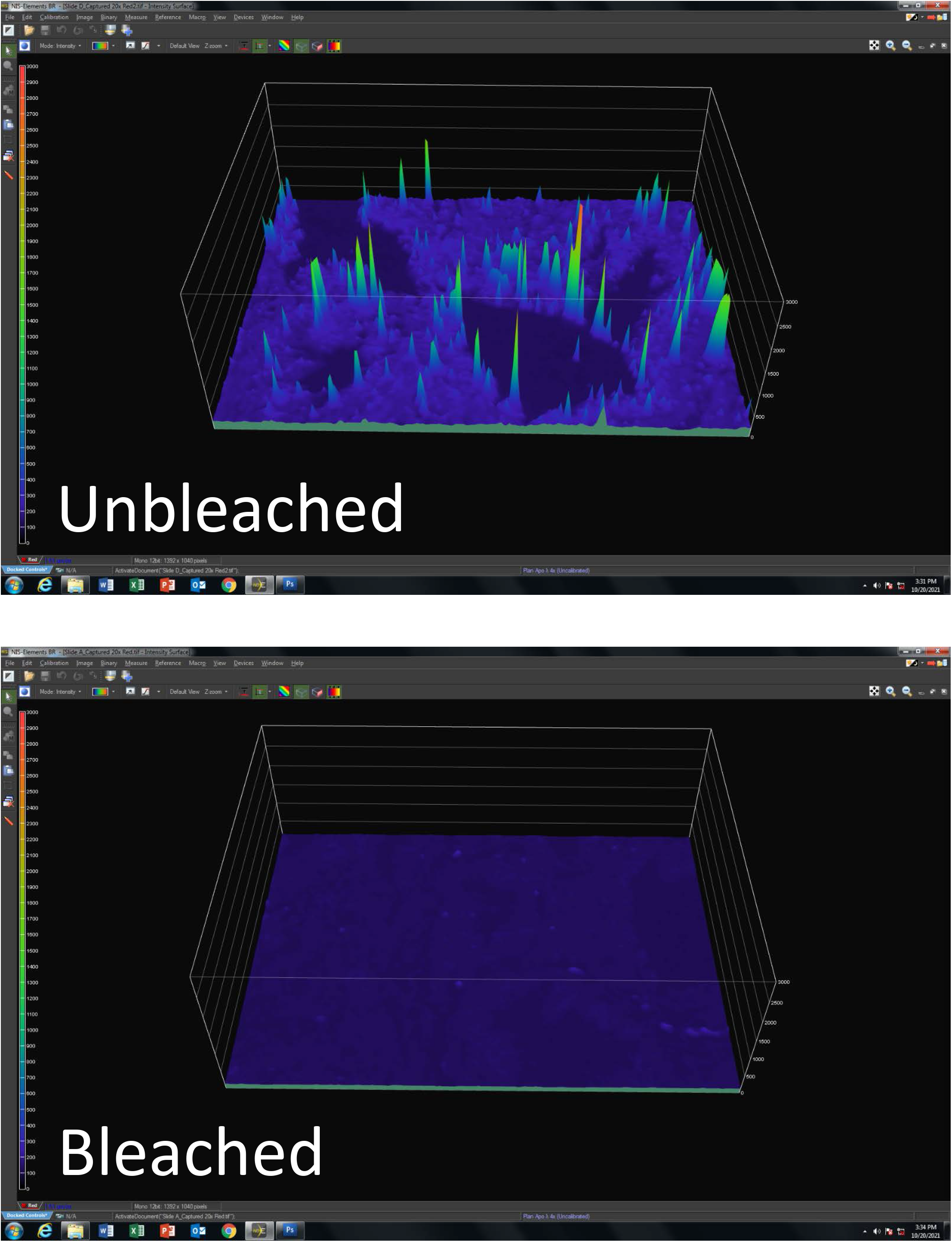
3D contour representations of reduction in autofluorescence by photochemical bleaching in FFPE prostate sections. Shown here are fluorescence contour plots comparing an unbleached FFPE tissue section (upper panels) and a bleached adjacent section (lower panels). A) Green channel; B) Red channel. For each color channel, exposure times used in image capture were identical for unbleached and bleached slides and the plots for each color are scaled to the same minimum and maximum values on the z-axis. All images were captured using a 20X objective.

In addition to following the published photochemical bleaching protocol, we also explored variations on the protocol including, varying the number of treatment cycles between one to four, treating the slides with hydrogen peroxide solution only, or with LED light exposure only (slides immersed in PBS), as well as substituting bright afternoon sunlight as the light source in the standard protocol. When comparing different numbers of treatment cycles, we found that a single cycle was markedly less effective than two cycles in reducing autofluorescence, while three cycles exhibited only minor additional autofluorescence suppression, compared to the standard two cycles, and no noticeable additional suppression with the addition of one more cycle (four cycles total, data not shown). Although we did observe noticeable diminution in autofluorescence for slides treated with either bleaching solution alone or LED light exposure alone, neither single treatment produced the dramatic decreases seen when both are used in combination (data not shown). Interestingly, conducting two cycles of bleaching and substituting bright sunlight exposure in place of the LED light source resulted in nearly complete elimination of autofluorescence in the tissue (data not shown).

To test the compatibility of the published protocol with subsequent immunofluorescent staining, we conducted immunofluorescence on serial sections of fixed prostate tissues either unbleached or bleached with two hydrogen peroxide/LED light cycles, as per protocol. As seen in **Figure 4A**, in the unbleached sample (top image), specific identification of the three true CD4-positive T cells stained in green (red arrows), is difficult to observe due to significant interference from background endogenous fluorescence; whereas, bleaching prior to immunostaining facilitates identification of the five CD4-positive T cells, due to the increased signal-to-noise obtained by suppression of the autofluorescence in this adjacent section. **Figure 4B** shows results in a section from a separate fixed prostate tissue specimen, dual-stained for both CD4-positive (green) and CD8-positive (red) T cells. As observed in the unbleached tissue section (top panel), this case harbors intense autofluorescence in both the green and red channels, making it extremely difficult to clearly identify the presence of either type of lymphocyte by their respective immunostains. In contrast, an adjacent section from the same tissue block that was bleached prior to multiple IF staining (lower panel) demonstrates dramatically improved signal-to-noise achieved, thus facilitating the identification of both types of lymphocytes present in the sample. This case further highlights the benefit that bleaching provides in cases where the IF signals are weak-to-moderate (here, white arrows indicate examples of CD4-positive T cells moderately stained in green). Other than the specimen shown in **Figure 1A–C**, for all cases where specimens are directly compared, image sets were collected in the same imaging sessions, using identical exposure times for each separate color channel, merged into single multicolor composite images and treated identically in post-processing. For the slides shown in **Figure 4**, cell nuclei were stained in blue with the fluorescent DNA-binding dye, DAPI. No significant reductions in DAPI fluorescence were observed when unbleached and bleached samples were compared (data not shown).

**Figure 4.**
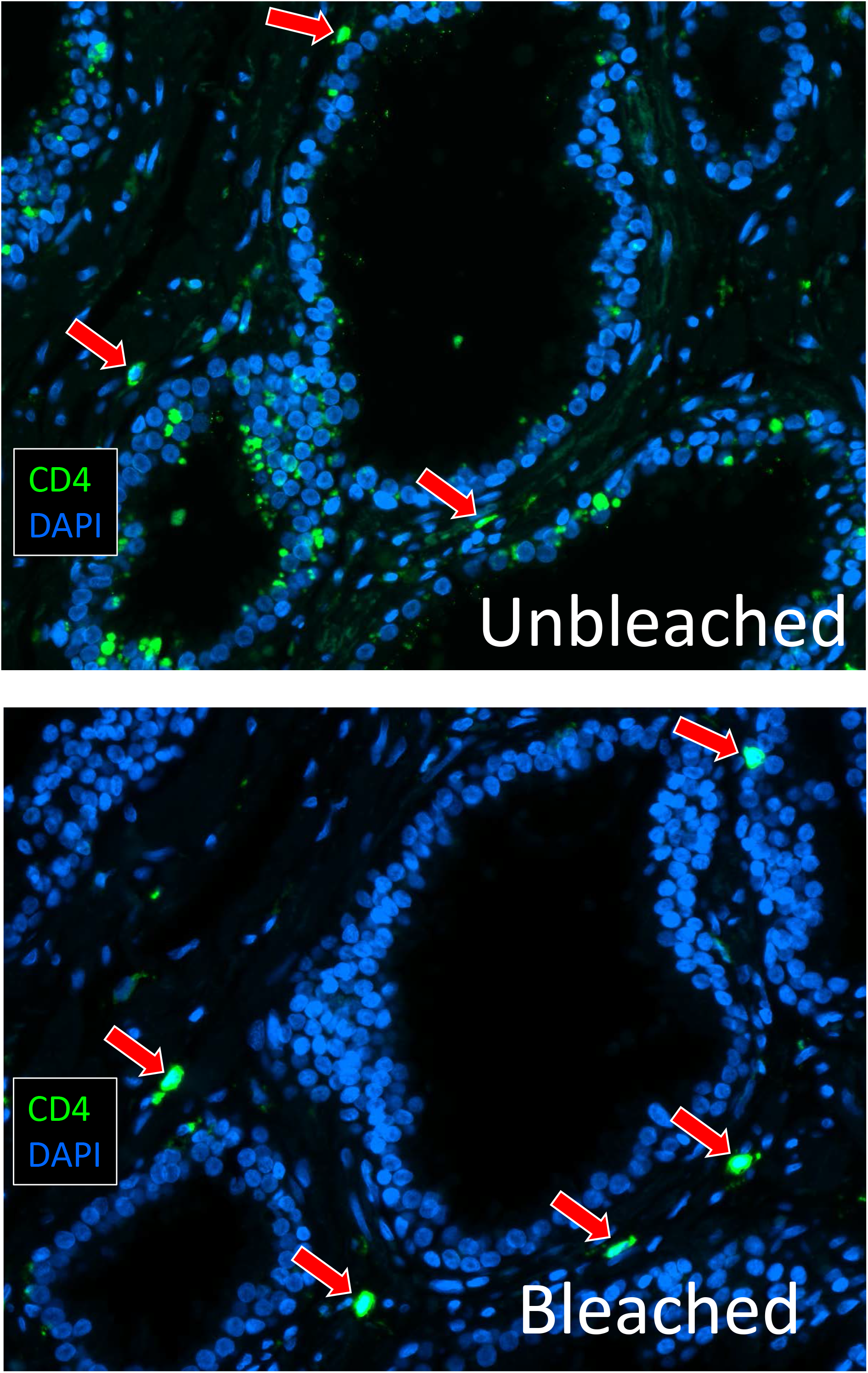
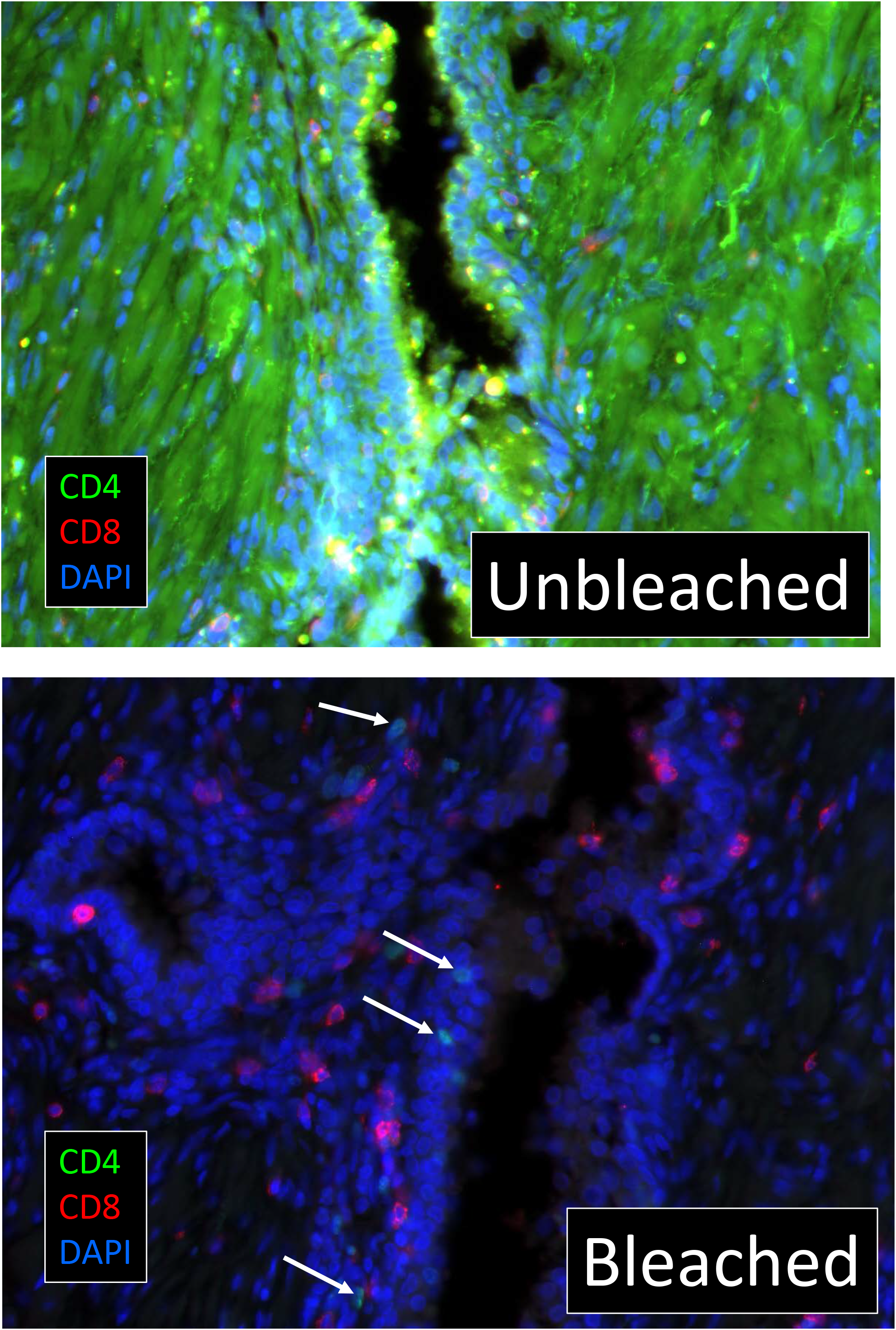
Autofluorescence photochemical bleaching is compatible with immunofluorescence and facilitates identification of fluorescently stained target cells in FFPE prostate sections. A) Adjacent sections, either unbleached (upper) or bleached (lower), that were then stained with an anti-CD4 antibody and detected by green immunofluorescence. Red arrows indicate all true CD4-positive cells, as determined by co-localization with cell nuclei stained in blue with DAPI. B) Adjacent sections, either unbleached (upper) or bleached (lower), that were then stained using dual-immunofluorescence with an anti-CD4 antibody (green) and an anti-CD8 antibody (red). White arrows in the lower image indicate representative CD4-positive cells with weak-to-moderate immunofluorescence staining that are nonetheless clearly visible.

## Discussion and Conclusions

In biomedical studies employing multiplex IF, endogenous tissue autofluorescence often interferes with detection of the desired antibody-derived fluorescent signals by decreasing signal-to-noise, in the worst cases preventing detection of the IF signals altogether. We have observed this to be especially problematic when performing cell phenotyping via multiplex IF on fixed human prostate cancer tissue sections. Our laboratory conducts molecular patho-epidemiologic studies in prostate cancer requiring multiplex IF staining of hundreds of FFPE prostate specimens, sourced from large clinical and epidemiologic cohorts. The tissue samples in these cohorts are typically FFPE blocks sourced from surgical pathology archives, and have been in storage for many years to decades. Prostate cancer largely afflicts older men, thus these tissues have accumulated substantial amounts of age-related autofluorescent material over the many decades of the patients’ lives. An effective method for reducing or eliminating autofluorescence in archival FFPE human prostate tissues would be of great benefit. Recently, Du et al. described a method for reducing autofluorescence in FFPE tissue specimens and is compatible with IF (25). Here, we demonstrate that this method is highly effective at decreasing the endogenous autofluorescent background in sections cut from archival FFPE prostate tissue blocks. We find that the published method, featuring two sequential 45 minute rounds of bleaching via slide immersion in alkaline hydrogen peroxide solution with simultaneous exposure to a bright white LED light source, reduces the brightest autofluorescence signals by 80%, in multiple wavelengths across the visible spectrum that are commonly used in IF. Additionally, we find that this effect is durable for at least one month (the longest time point assessed). By varying the number of bleaching cycles, we determined that a single cycle is markedly less effective at reducing autofluorescence, whereas, three or more cycles provide little added benefit compared to the two cycles recommended in the published protocol. Interestingly, we found that substituting bright mid-day sunlight in place of the artificial LED light source resulted in a near complete suppression of all autofluorescence in the tissue section. While obviously difficult to routinely and reproducibly use sunlight in practice, this finding strongly suggests that artificial light sources that more closely replicate the spectrum and/or intensity of natural sunlight may provide further decreases in autofluorescence, and thus warrants further study.

As previously demonstrated by Du et al for other tissue types, we demonstrate that this photochemical autofluorescence bleaching method is compatible with subsequent multiplex IF staining, as well as nuclear staining by the DNA-specific dye, DAPI. Importantly, we show that bleaching FFPE prostate tissue sections prior to IF markedly increases signal-to-noise, thus facilitating detection of the desired IF signals, in multiple color channels, and is particularly helpful where the autofluorescence is very intense and/or the IF signals are weak.

## Acknowledgements

This work was funded through the following grants and awards: R01 CA255349; W81XWH-20-1-0264; P30 CA006973.

## References

1. Manos PD, Ratanasirintrawoot S, Loewer S, Daley GQ, Schlaeger TM. Live-cell immunofluorescence staining of human pluripotent stem cells. Curr Protoc Stem Cell Biol. 2011 Dec; Chapter 1:Unit 1C.12. PMID: 22135082.

2. Stack EC, Wang C, Roman KA, Hoyt CC. Multiplexed immunohistochemistry, imaging, and quantitation: a review, with an assessment of Tyramide signal amplification, multispectral imaging and multiplex analysis. Methods. 2014 Nov;70(1):46–58. PMID: 25242720.

3. Abuduwali N, Lossdörfer S, Winter J, Wolf M, Götz W, Jäger A. Autofluorescent characteristics of human periodontal ligament cells in vitro. Ann Anat. 2013 Oct;195(5):449–54. PMID: 23706696.

4. Yang Y, Honaramooz A. Characterization and quenching of autofluorescence in piglet testis tissue and cells. Anat Res Int. 2012;2012:820120. PMID: 22970375; PMCID: PMC3437279.

5. Georgakoudi I, Jacobson BC, Müller MG, Sheets EE, Badizadegan K, Carr-Locke DL, Crum CP, Boone CW, Dasari RR, Van Dam J, Feld MS. NAD(P)H and collagen as in vivo quantitative fluorescent biomarkers of epithelial precancerous changes. Cancer Res. 2002 Feb 1;62(3):682–7. PMID: 11830520.

6. Menter JM. Temperature dependence of collagen fluorescence. Photochem Photobiol Sci. 2006 Apr;5(4):403–10. PMID: 16583021.

7. Zipfel WR, Williams RM, Christie R, Nikitin AY, Hyman BT, Webb WW. Live tissue intrinsic emission microscopy using multiphoton-excited native fluorescence and second harmonic generation. Proc Natl Acad Sci U S A. 2003 Jun 10;100(12):7075–80. PMID: 12756303; PMCID: PMC165832.

8. Gallas, J.M. and Eisner, M. Fluorescence of melanin-dependence upon excitation wavelength and concentration. Photochemistry and Photobiology, 1987; 45: 595–600.

9. Monici M. Cell and tissue autofluorescence research and diagnostic applications. Biotechnol Annu Rev. 2005;11:227–56. PMID: 16216779.

10. Whittington NC, Wray S. Suppression of Red Blood Cell Autofluorescence for Immunocytochemistry on Fixed Embryonic Mouse Tissue. Curr Protoc Neurosci. 2017 Oct 23;81:2.28.1–2.28.12. PMID: 29058770; PMCID: PMC5657453.

11. Willingham MC. An alternative fixation-processing method for preembedding ultrastructural immunocytochemistry of cytoplasmic antigens: the GBS (glutaraldehyde-borohydride-saponin) procedure. J Histochem Cytochem. 1983 Jun;31(6):791–8. PMID: 6404984.

12. Viegas MS, Martins TC, Seco F, do Carmo A. An improved and cost-effective methodology for the reduction of autofluorescence in direct immunofluorescence studies on formalin-fixed paraffin-embedded tissues. Eur J Histochem. 2007 Jan-Mar;51(1):59–66. PMID: 17548270.

13. Schönenbrücher H, Adhikary R, Mukherjee P, Casey TA, Rasmussen MA, Maistrovich FD, Hamir AN, Kehrli ME Jr, Richt JA, Petrich JW. Fluorescence-based method, exploiting lipofuscin, for real-time detection of central nervous system tissues on bovine carcasses. J Agric Food Chem. 2008 Aug 13;56(15):6220–6. PMID: 18620407.

14. Goldin A, Beckman JA, Schmidt AM, Creager MA. Advanced glycation end products: sparking the development of diabetic vascular injury. Circulation. 2006 Aug 8;114(6):597–605. PMID: 16894049.

15. Clancy B, Cauller LJ. Reduction of background autofluorescence in brain sections following immersion in sodium borohydride. J Neurosci Methods. 1998 Sep 1;83(2):97–102. PMID: 9765122.

16. Schnell SA, Staines WA, Wessendorf MW. Reduction of lipofuscin-like autofluorescence in fluorescently labeled tissue. J Histochem Cytochem. 1999 Jun;47(6):719–30. PMID: 10330448.

17. Yang J, Yang F, Campos LS et al. Quenching autofluorescence in tissue immunofluorescence Wellcome Open Res 2017, 2:79 (https://doi.org/10.12688/wellcomeopenres.12251.1)

18. Davis AS, Richter A, Becker S, Moyer JE, Sandouk A, Skinner J, Taubenberger JK. Characterizing and Diminishing Autofluorescence in Formalin-fixed Paraffin-embedded Human Respiratory Tissue. J Histochem Cytochem. 2014 Jun;62(6):405–423. PMID: 24722432; PMCID: PMC4174629.

19. Neumann M, Gabel D. Simple method for reduction of autofluorescence in fluorescence microscopy. J Histochem Cytochem. 2002 Mar;50(3):437–9. PMID: 11850446.

20. Levenson, R.M. and Mansfield, J.R. (2006), Multispectral imaging in biology and medicine: Slices of life. Cytometry, 69A: 748–758.

21. Mansfield JR. Multispectral imaging: a review of its technical aspects and applications in anatomic pathology. Vet Pathol. 2014 Jan;51(1):185–210. PMID: 24129898.

22. Berry S, Giraldo NA, Green BF, Cottrell TR, Stein JE, Engle EL, Xu H, Ogurtsova A, Roberts C, Wang D, Nguyen P, Zhu Q, Soto-Diaz S, Loyola J, Sander IB, Wong PF, Jessel S, Doyle J, Signer D, Wilton R, Roskes JS, Eminizer M, Park S, Sunshine JC, Jaffee EM, Baras A, De Marzo AM, Topalian SL, Kluger H, Cope L, Lipson EJ, Danilova L, Anders RA, Rimm DL, Pardoll DM, Szalay AS, Taube JM. Analysis of multispectral imaging with the AstroPath platform informs efficacy of PD-1 blockade. Science. 2021 Jun 11;372(6547):eaba2609. doi: 10.1126/science.aba2609. PMID: 34112666.

23. Christian, J., Lamm, T., Morrow, J. et al. Corpora amylacea in adenocarcinoma of the prostate: incidence and histology within needle core biopsies. Mod Pathol 18, 36–39 (2005).

24. Brennick, Jeoffry B. M.D.; O’Connell, John X. M.B. B.ch. F.R.C.P.C.; Dickersin, G. Richard M.D., Ben Z. Pilch, M.D.; Young, Robert H. M.D. M.R.c.Path. Lipofuscin Pigmentation (So-Called “Melanosis”) of the Prostate, The American Journal of Surgical Pathology: May 1994 - Volume 18 - Issue 5 - p 446–454.

25. Du, Z., Lin, JR., Rashid, R. et al. Qualifying antibodies for image-based immune profiling and multiplexed tissue imaging. Nat Protoc. 2019 Oct;14(10):2900–2930 (2019). PMID: 31534232 PMCID: PMC6959005

